# YAP levels regulate anteroposterior elongation of hESC-derived gastruloids

**DOI:** 10.64898/2025.12.01.691265

**Authors:** Elizabeth Abraham, Thomas Roule, Olivia Mae Pericak, Mikel Zubillaga, Naiara Akizu, Conchi Estaras

**Affiliations:** Department of Cardiovascular Sciences, Aging + Cardiovascular Discovery Center, Temple University, Lewis Katz School of Medicine, Philadelphia, PA, USA; Cancer Epigenetics Institute, Fox Chase Cancer Center, Philadelphia, Pennsylvania, USA; Raymond G. Perelman Center for Cellular and Molecular Therapeutics, The Children’s Hospital of Philadelphia, Philadelphia, PA, USA

**Keywords:** gastrulation, YAP1, 3D gastruloids, symmetry breaking, hESCs, Hippo pathway

## Abstract

Human embryonic stem cells (hESCs) can self-organize into anteroposterior patterned three-dimensional structures, characterized by polarized expression of CDX2 and GATA6, known as *human 3D gastruloids (h3D-gastruloids)*. This patterning emerges through hESC-intrinsic mechanisms that remain poorly understood. Here, we demonstrate that the formation and elongation of *h3D*-gastruloids is modulated by the activity levels of the Hippo pathway effector YAP. Using complementary chemical and genetic perturbations, we show that elevated nuclear YAP activity inhibits gastruloid elongation, whereas YAP inhibition enhances axial elongation relative to controls. Single-cell RNA sequencing (scRNA-seq) analyses reveal that YAP activation disrupts the establishment of distinct anterior GATA6 and posterior CDX2 poles. Conversely, YAP inhibition promotes clearer segregation of these domains, facilitating symmetry breaking and elongation. Finally, we developed a high-throughput platform for *h3D*-gastruloid generation, enabling the production of over 500 gastruloids per well. Together, our findings uncover a role for YAP in orchestrating the three-dimensional organization of human gastruloids.

## INTRODUCTION

Gastrulation is a milestone event in early embryonic development that originates the three germ layers and the body plan^1,2^. Errors in this process are a major cause of infertility, miscarriage, and birth defects^3–6^, all of which impose a significant burden to society and the health system. Although decades of research using model organisms have uncovered fundamental mechanisms of early embryonic development, elucidating human disease mechanisms will require unfolding the complexity of human embryonic development.

Human embryonic stem cell-derived 3D-gastruloids (h3D-gastruloids) recapitulate key features of human gastrulas^7^. These h3D-gastruloids exhibit organized cell populations and structures that resemble the mammalian primitive streak and tailbud. Unlike embryoid bodies, h3D-gastruloids can break the radial symmetry of hESC spheres, establishing compartmentalized signaling and gene expression components along an anteroposterior axis. This capability aligns emerging cell populations along the axis, initiating an elongation process that resembles core elements of the body plan^1,7^. This body plan, or blueprint, encodes essential lineage-determination cues and spatial organization across anterior-posterior, mediolateral, and dorsoventral axes^1,7^. Thus, symmetry breaking and elongation are essential processes in body plan formation, as captured in h3D-gastruloids. Notably, the mechanisms controlling these processes are only partially known^8–14^.

Long-standing in vivo studies in mouse embryos have shown that gastrulation and the establishment of the anteroposterior axis are regulated by a gradient of Nodal:Smad2/3 and Wnt:β-catenin signaling shaped by extraembryonic signals. At the onset of gastrulation, BMP signals from the extraembryonic ectoderm induce Nodal and Wnt activities on the posterior epiblast to specify the Primitive Streak (PS)^15^. Inversely, the extraembryonic anterior visceral endoderm (AVE) sends inhibitory Wnt and Nodal signals to the anterior epiblast, promoting ectoderm differentiation^16,17^. However, h3D-gastruloids in vitro organize in the absence of extraembryonic tissues.

During h3D-gastruloids formation, hESC-spheres undergo significant morphological and signaling changes after incubation with Wnt agonists. These changes result in the development of a posterior pole, characterized by the expression of CDX2, T/BRACHYURY, and active Wnt and Nodal signals. On the other side of the structure, an anterior pole forms, populated by anterior mesoderm, including derivatives of cardiac lineages, and broadly marked by GATA6 expression and active BMP signaling^7^. The asymmetric signaling gradient is thought to guide a morphogenetic process where cells intercalate and elongate along an axis, causing the gastruloid to lengthen while narrowing in width. However, few mechanisms are known to contribute to the organization of the h3D-gastruloid signaling and formation^8–14^, and we are thus far from fully comprehending how the antero-posterior axis is generated in response to spatially uniform cues.

The Hippo: YAP signaling pathway is essential for development and regulates various processes, including cell proliferation and differentiation^18–21^. When the Hippo kinases are inactive, YAP translocates into the nucleus, where it binds to TEAD transcription factors to regulate gene expression^21^. Our work, along with that of others, has shown that YAP modulates gastrulation signaling genes and suppresses mesoderm differentiation during hESCs-directed differentiation and in 2D gastruloid models^22,23^. However, the role of YAP in regulating gastrulation within a three-dimensional context remains unexplored.

Hence, in this study, we investigated the contribution of YAP levels to the formation of the h3D-gastruloids. By combining chemical and genetic approaches, we found that YAP depletion results in robust elongation, yielding longer h3D-gastruloids than controls. On the contrary, forcing nuclear YAP retention by inhibition of the Hippo kinases abolishes elongation, resulting in the formation of hESC-spheres. Single-cell RNAseq analysis revealed that YAP activation impairs the establishment of distinct anterior and posterior poles and increases the proportion of double-positive CDX2⁺/GATA6⁺ cells relative to controls. Instead, YAP inhibition accelerates the differentiation of epiblast-like cells toward mesodermal lineages, which correlates with robust and elongated gastruloids formation. Overall, our studies reveal that YAP levels regulate anteroposterior elongation of h3D-gastruloids.

## RESULTS

### Efficient generation of hESC-derived 3D gastruloids in microwells

In this study, we analyze a frontline model that recapitulates human gastrulation in 3D^7^ to investigate novel mechanisms controlling the spatiotemporal organization of gene expression. We adapted the h3D-gastruloid generation method to use on microwell plates (AggreWell^TM^). This approach, tested in multiple hESC and iPSCs lines, enabled simultaneous induction of large numbers of gastruloids per condition, reducing the bias and increasing the accuracy of analysis (**Figure 1A**). Following pretreatment with the Wnt activator CHIR99021, hESC colonies are dissociated and cells are seeded on the microwells, where they aggregate in spherical hESC-bodies that break symmetry in the next 24h, followed by robust anteroposterior elongation growth over the course of 72h (**Figure 1B-C**). As expected, the three germinal layers were visualized (**Figure 1D**) and the organization of posterior-to-anterior cell populations was confirmed by the complementary patterns of CDX2 and GATA6 markers^7^ (**Figure S1A**).

**Figure 1:**
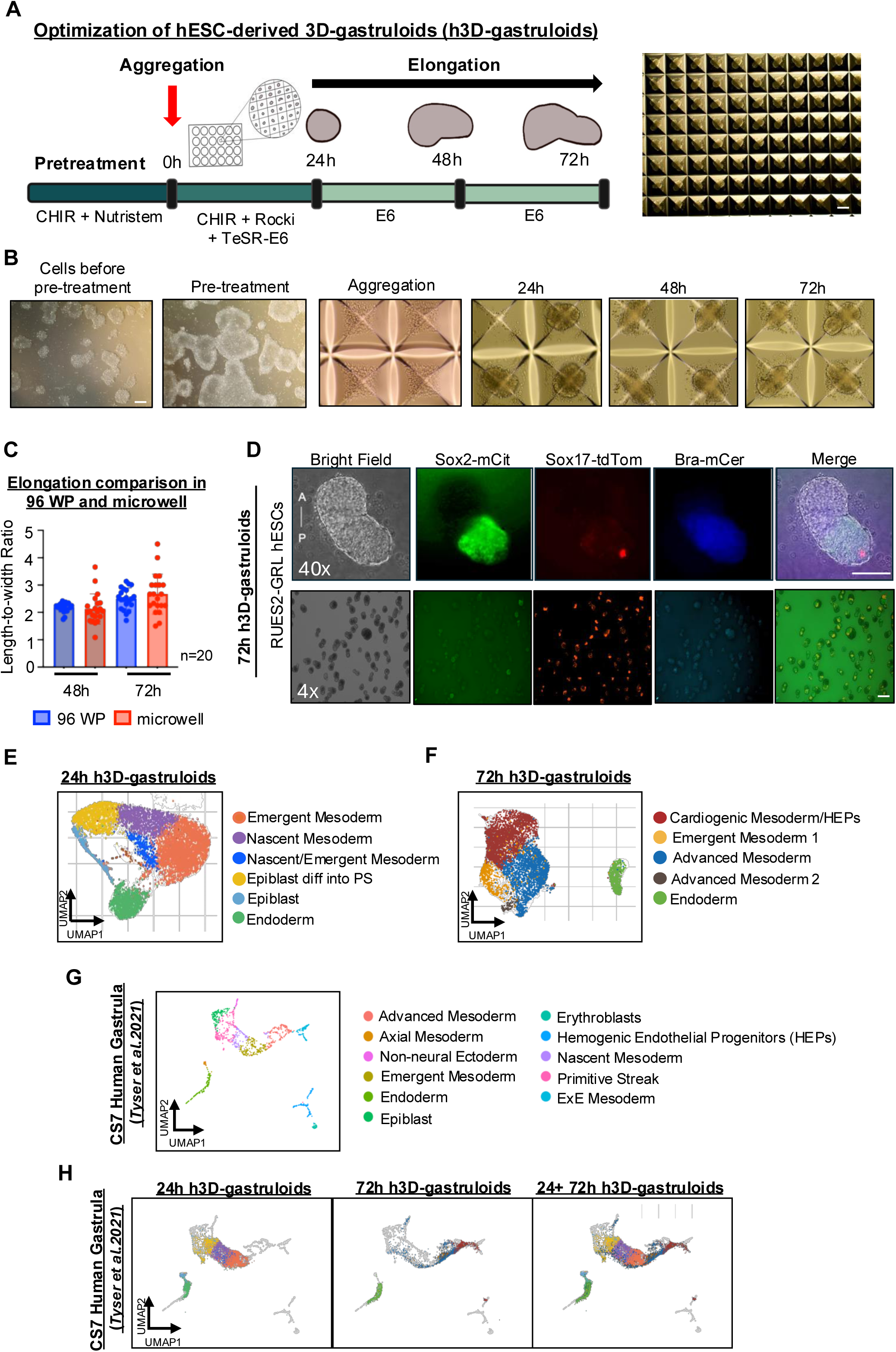
Generation and characterization of hESC-derived 3D gastruloids using a microwell system. (A) Schematic overview of the workflow for generating human 3D gastruloids (h3D-gastruloids) from human embryonic stem cells (hESCs) using a microwell-based aggregation system. Representative bright-field image of h3D-gastruloids growing in microwells. Scale bar, 125 µm. (B) Bright-field images of RUES2 hESCs from pre-treatment, aggregation, and culture through 72 hours. Scale bar, 125 µm. (C) Quantification of elongation based on length-to-width ratio comparing gastruloids generated in U-bottom 96-well plates versus microwells. Quantification was performed using the method described in Moris et al., 2020. Data was collected from two separate experiments and 10 gastruloids were counted each time, n=20. Data are presented as mean ± SEM. (D) Fluorescence images of 72-hour RUES2-GRL h3D-gastruloids showing posterior localization of the three germ layers using lineage-specific reporters (mCit = mCitrine, tdTom = tdTomato, mCer = mCerulean). Two magnifications are shown. *Scale bars, 125 µm*. (E) UMAP visualization of single-cell RNAseq from 24-hour h3D-gastruloids with annotated cell populations (F) UMAP representation of single-cell RNAseq from 72h h3D-gastruloids with annotated populations (G) UMAP representation of the CS7 human gastrula single-cell transcriptomic dataset published by Tyser *et al*., 2021. (H) Projection of 24- and 72-hour h3D-gastruloid single-cell transcriptomes onto the human gastrula reference atlas from Tyser *et al.*, 2021.

We performed single-cell RNA sequencing (scRNA-seq) analysis at 24 and 72 hours (see Methods for details). At 24 hours, the gastruloids recapitulated key early developmental clusters, including epiblast-like cells (POU5F1⁺/SOX2⁺), primitive streak (TBXT⁺), endoderm (FOXA2⁺/SOX17⁺), and nascent (TBXT^low^/EOMES⁺/MESP1⁺) and emergent mesoderm populations (EOMES⁺/CER1⁺/GSC⁺) (**Figure 1E and Figure S1B**). By 72 hours, the epiblast-like cluster was no longer present, having fully transitioned into differentiated cell types (**Figure 1F**). Endoderm clusters persisted, along with nascent and emergent mesoderm populations (MESP1⁺). In addition, subsets of mesodermal cells showed further maturation into advanced lineages expressing markers of cardiovascular mesoderm (MESP1^low^/KDR⁺/GATA4⁺), with a fraction within this cluster clearly progressing toward hematopoietic progenitors (TAL1⁺) (**Figure 1F** and **Figure S1C**).

To validate that h3D-gastruloids faithfully model early human development, we mapped their transcriptomes onto a CS7 human gastrula reference (**Figure 1G**) and computed a score reflecting global transcriptional similarity (see Materials and Methods)^24^. All major populations from both 24h and 72h gastruloids exhibited strong similarity to corresponding CS7 gastrula cell types (**Figure 1H**). As expected, 24h gastruloid populations aligned preferentially with earlier gastrula states, including epiblast and primitive streak clusters (**Figure S1D**). By contrast, the 72h cardiogenic mesoderm population mapped to CS7 advanced mesoderm and hematopoietic progenitor clusters, consistent with further lineage maturation (**Figure S1E**).

Altogether, these analyses demonstrate that our scale-up platform generates h3D-gastruloids that recapitulate the cellular diversity and developmental progression of the human gastrula (see Methods for details).

### YAP levels regulate h3D-gastruloid formation

We have previously shown that the transcriptional repressor activity of YAP1 regulates mesodermal differentiation during 2D-hESC differentiation and in mouse models^25,26^. To assess whether YAP activity affects h3D-gastruloid formation, we treated the gastruloids at the time of aggregation with the small molecules Dasatinib (YAPi) or XMU-MP-1 (YAPa), which inhibit or promote nuclear YAP accumulation, respectively^27,28^ (**Figure 2A**). Dasatinib works by inhibiting YES1 and Src-family kinases, forcing YAP accumulation in the cytosol, while XMU-MP-1, forces nuclear retention of YAP by inhibiting the MST1/2 kinases^27,28^. To confirm the effectiveness of these treatments, we validated their impact on nuclear YAP accumulation by qPCR and western blot, showing that both compounds effectively modulate YAP distribution (**Figure 2B and Figure S2A-B).** Furthermore, ChIP-qPCR at the known YAP target locus, CTGF, demonstrated that YAP binding is altered by both activator and inhibitor treatment (**Figure 2C**). h3D-gastruloids treated with the Dasatinib (YAPi) exhibit significantly enhanced elongation compared to controls (**Figure 2D-E** and **Figure S2C)**. In contrast, treatment with XMU-MP-1 (**YAPa**) led to the formation of rounded gastruloids, unable to break radial symmetry (**Figure 2D-E** and **Figure S2C**).

**Figure 2:**
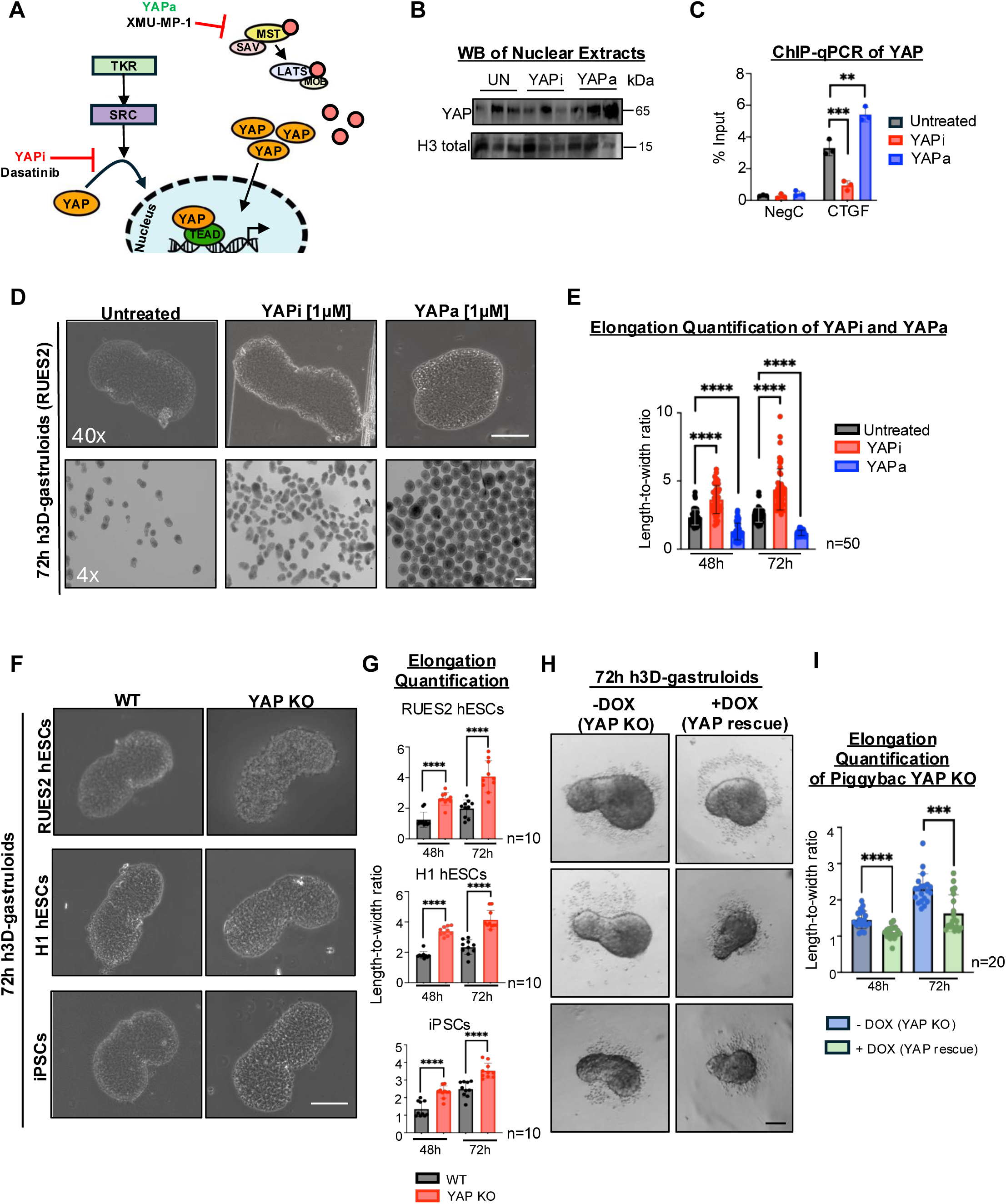
YAP levels regulate elongation of human 3D gastruloids. (A) Scheme showing the mechanism of action of the small molecules Dasatinib (YAP inhibitor; YAPi) and XMU-MP-1 (YAP activator; YAPa) in modulating YAP activity. (B) Western blot analysis of nuclear extracts from WT H1 hESCs treated with 1 µM YAPi or YAPa for 48 hours, probed for YAP and Histone H3. Experiment shown was performed once with three technical replicates. (C) ChIP-qPCR analysis of YAP in WT hESCs treated with YAPi and YAPa for 48 hours. Genomic regions tested are indicated in the graph. Data are presented as mean ± SEM. Experiment was performed once with three technical replicates. Statistical analysis: one-way ANOVA (Kruskal-Wallis test) ^∗∗^*p* < 0.01, ^∗∗∗^*p <* 0.001. NegC: negative control region (D) Representative bright-field images of 72-hour RUES2-derived h3D-gastruloids untreated or treated with YAPi or YAPa at the indicated concentrations during aggregation. Two magnifications are shown. Scale bar, 125 µm. (E) Quantification of gastruloid elongation (length-to-width ratio) for RUES2-derived h3D-gastruloids untreated or treated with the indicated molecules. Quantification was performed as described in Moris *et al.* (2020). Data are shown as mean ± SEM. Data represent three independent experiments (15–20 gastruloids per experiment; total *n* = 50). Statistical analysis: one-way ANOVA (Kruskal–Wallis). ****p < 0.0001. (F) Representative bright-field images of WT and YAP KO gastruloids generated from multiple pluripotent stem cell lines (RUES2, H1, and iPSCs) and cultured for 72 hours. Scale bar, 125 µm. (G) Quantification of elongation (length-to-width ratio) in WT and YAP KO h3D-gastruloids at 48 and 72 hours. Data are shown as mean ± SEM. Data represent two independent experiments (5 gastruloids per experiment; total *n* = 10). Statistical analysis: Student’s *t*-test. ****p < 0.0001. (H) Representative bright-field images of 72-hour h3D-gastruloids derived from Piggybac YAP KO hESCs. In this inducible system, Doxycycline restores YAP expression (+DOX, 24h). Doxycycline treatment reduces gastruloid elongation compared with untreated YAP KO control (-DOX). Scale bar, 200 μm. (I) Quantification of elongation (length-to-width ratio) for gastruloids shown in **H**. Data are shown as mean ± SEM. Data represent two independent experiments (10 gastruloids per experiment; total *n* = 20). Statistical analysis: Student’s *t*-test. ****p < 0.0001.

To further examine the contribution of YAP to the 3D organization of the gastruloids, we applied CRISPR NHEJ to knockout the YAP gene in the RUES2 hESC line (**Figure S2D**). YAP deletion in the clonal KO line did not compromise the ability of hESCs to form pluripotent colonies, in line with what we reported in H1 background^29^ (**Figure S2E-F**). Thus, we examined the formation of h3D gastruloids in the WT and the YAP KO RUES2 hESC lines. We observed similar gastruloid formation efficiency in WT and YAP KO cells. However, YAP KO aggregates develop into longer gastruloids over time, suggesting more efficient axial elongation, compared to controls (**Figure 2F-G** and **Figure S2F**). Importantly, similar results were obtained in H1 hESCs and iPSCs lines, indicating that YAP deletion’s phenotype is conserved across distinct human genetic backgrounds (**Figure 2F-G** and **Figure S2G)**. Furthermore, we used a Doxycycline-inducible PiggyBac cell line to rescue YAP levels in the YAP KO H1 background^29^. YAP rescue led to shorter, spheroid-like gastruloids, compared to YAP KO (**Figure 2H-I**). Overall, these findings demonstrate that nuclear accumulation of YAP is incompatible with elongation, whereas its inhibition promotes gastruloid development along the anteroposterior axis.

### Cell population analysis of the h3D-gastruloids

To investigate the cell populations regulated by YAP during gastruloid formation, we performed scRNA-seq analysis of 24-hour and 72-hour gastruloids treated with YAPi or YAPa, and compared them to untreated controls. For each condition, around 1000 gastruloids were pooled and sequenced, yielding high-quality data. More than 4,000 cells per condition passed a stringent filtering criterion, enabling efficient clustering and robust identification of distinct populations (see Methods for details). Treatment with YAPi resulted in a robust downregulation of several established YAP-target genes, including CCN1 and CCN2, members of the IGFBP family (IGFBP2 and IGFBP5)^30^, and Hippo pathway regulators such as AJUBA and AMOT (**Figure S3A**). Conversely, many of these YAP-target signature genes were upregulated in YAPa-treated gastruloids, as expected, further confirming the effectiveness of the small-molecule treatments (**Figure S3A**).

### YAP inhibition promotes differentiation

Notably, the integrative analysis revealed that YAPi-treated cells were distributed across annotated clusters in proportions distinct from those observed in controls (**Figure 3A-B**). Among these clusters, the epiblast-like cluster (hESCs-like) exhibited the most pronounced differences and a marked reduction in cell content at 24h (**Figure 3A-B**). This depletion could potentially result from increased cell death, a prolonged cell cycle, or an elevated differentiation rate. However, cell cycle analysis revealed no significant differences between control and YAP-inhibited gastruloids (**Figure S3B**). Similarly, differential gene expression analysis did not suggest an upregulation of apoptosis-related genes (**Table S1**). Taken together, these findings suggest that YAP inhibition enhances differentiation of the epiblast cluster toward mesodermal trajectories (**Figure 3C-D**), consistent with findings in the 2D differentiation systems and in vivo^25,26^.

**Figure 3:**
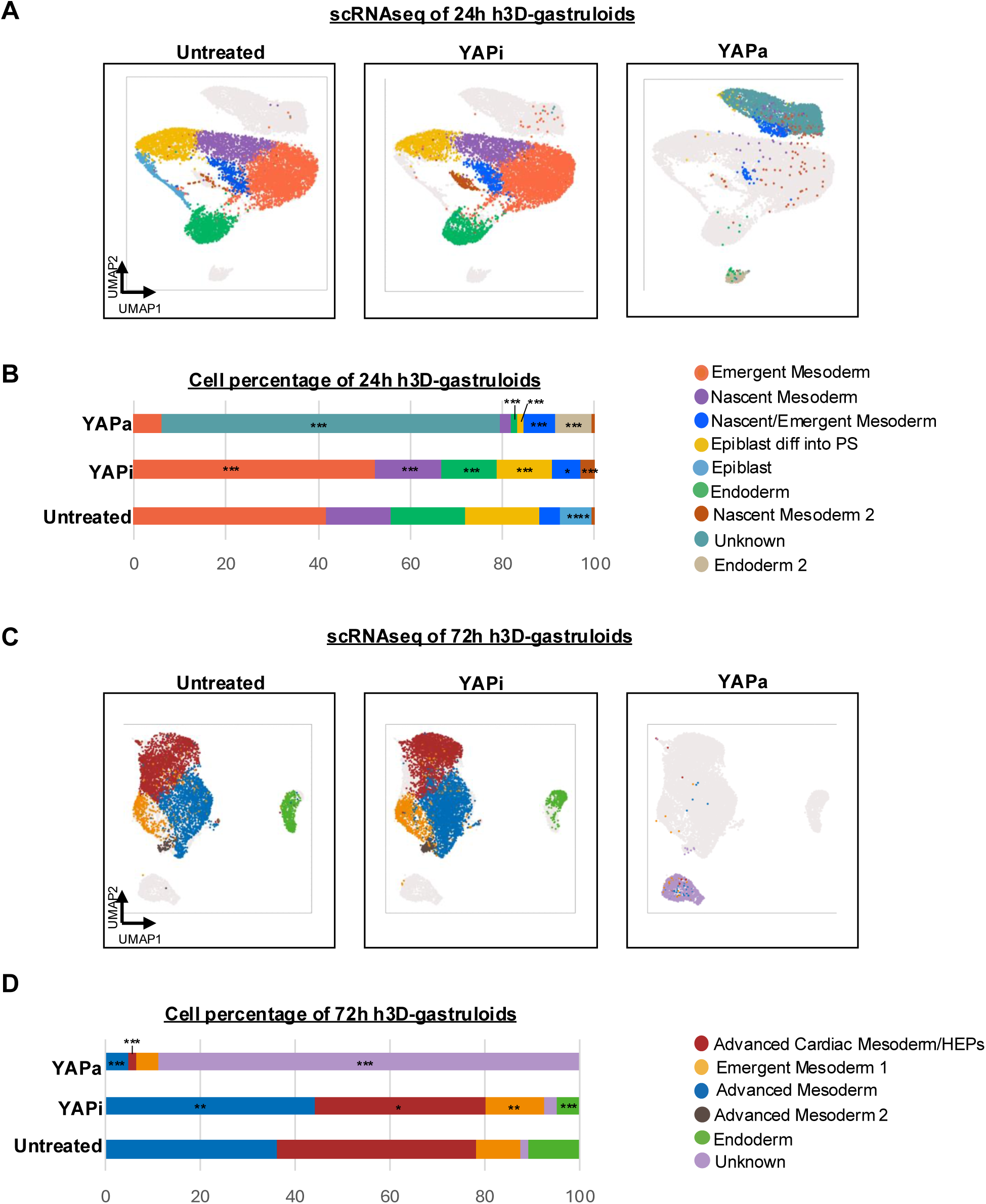
Effects of YAPi and YAPa treatment on cell populations in h3D-gastruloids. (A) UMAP projections of single-cell RNA sequencing from 24-hour h3D-gastruloids showing annotated cell populations for the untreated, YAP inhibitor (YAPi), and YAP activator (YAPa) conditions. (B) Proportion of cells within each annotated cluster across conditions at 24 hours. Data are derived from scRNA-seq. Statistical analysis: Chi-square test with Bonferroni correction. \**p* < 0.05*, **p < 0.01, ***p < 0.001*, \*\*\*\**p* < 0.0001 (C) UMAP projections of single-cell transcriptomes from 72-hour h3D-gastruloids showing annotated lineage populations for untreated, YAPi, and YAPa conditions. (D) Cell percentages of cells within each annotated cluster across conditions at 24 hours. Data are derived from scRNA-seq. Statistical analysis: Chi-square test with Bonferroni correction. \**p* < 0.05*, **p < 0.01, ***p < 0.001*, \*\*\*\**p* < 0.0001

### Aberrant lineage identities in YAP-activated gastruloids

YAPa-treated gastruloids at both 24-hour and 72-hour failed to align with any annotated clusters identified in control or YAPi conditions (**Figure 3A and Figure 3C**). Although subsets of YAPa cells expressed lineage-associated factors—including EOMES (mesoderm), TBXT (primitive streak), and epiblast-related genes such as NANOG and POU5F1—the absence of key defining markers prevented proper cluster assignment (**Figure S3C**). For example, POU5F1⁺ cells did not express SOX2,which prevented mapping to a bona fide epiblast cluster. A comparable disruption occurred in mesodermal populations. In control gastruloids, emergent mesoderm expressed CER1⁺ and EOMES⁺. Under YAP activation, EOMES remained detectable, but CER1 expression was almost entirely lost, indicating an incomplete or skewed mesodermal identity. Moreover, MESP1, normally restricted to nascent mesoderm became broadly expressed across most YAPa cells, reflecting an issue with lineage specificity. Cell-cycle analysis revealed no significant differences between YAPa and controls, consistent with a minimal role for YAP in controlling proliferation at this early developmental stage (**Figure S3B**).

Collectively, these findings indicate that forced nuclear YAP activity induces an aberrant transcriptional program that compromises lineage fidelity, blurs boundaries between early gastrulation identities, and prevents the formation of recognizable gastrula-like populations.

### Incomplete symmetry breaking in YAP-activated gastruloids

Defined anterior–posterior (A–P) domains are a hallmark of elongated h3D-gastruloids: anterior populations express GATA6, whereas posterior regions express CDX2 in largely non-overlapping territories (**Figure S4A**). To assess whether this organization is maintained under altered YAP activity, we quantified GATA6 and CDX2 expression in control, YAPa, and YAPi gastruloids. At 24 hours, control gastruloids contained ∼77.58% GATA6⁺ cells and ∼12.38% CDX2⁺ cells, with only ∼5.22% co-expressing both markers. YAPi gastruloids displayed a similar distribution (87.51% GATA6⁺, 12.84% CDX2⁺) and a comparable level of co-expression (∼7.74%) (**Figure 4A**). By 72 hour, however, YAPi gastruloids exhibited significantly reduced co-expression relative to controls (15.39% vs. 24.40%, adjusted *p* < 0.005), consistent with enhanced lineage segregation under YAP inhibition (**Figure 4B**). In sharp contrast, YAPa gastruloids showed an extensive co-expression pattern. At 24 hour, YAPa samples displayed widespread CDX2 expression (57.84% of cells), and a striking ∼48.18% of cells co-expressing both markers (vs. 5.2% in controls, *p* < 1×10⁻⁵). This effect persisted at 72 hour, where co-expression reached ∼57.18% (vs. 24.4% in controls, *p* < 1×10⁻⁵)(**Figure 4A-B**). Immunostaining of dissociated gastruloids corroborated this phenotype, revealing markedly elevated co-staining of GATA6 and CDX2 cells from YAPa gastruloids, compared to untreated samples (**Figure S4B**). Together, these findings reveal that YAPa gastruloids fail to form anterior or posterior domains, resulting in mixed lineage identities. To further examine anterior-posterior identity disruption, we extracted the top 100 expressed genes in CDX2⁺ cells in untreated gastruloids (“*CDX2+ cells signature genes*”) and compared their expression in YAPi and YAPa samples, both at 24 and 72 hours (**Figure 4C-D**). In 24-hour YAPa gastruloids, many of these CDX2 co-expressed genes were downregulated, including MEIS3 and WNT8A, whereas others, such as CDX2 itself and TBXT, were upregulated relative to untreated controls (**Figure 4C**). We next examined the expression of these 100 genes in GATA6⁺ cells in 24-hour gastruloids, where their expression is minimal in untreated gastruloids (**Figure 4C**). In YAPa gastruloids, many of these genes became ectopically expressed—including CDX2, TBXT, and WNT5B (**Figure 4C**), indicating that YAPa-GATA6⁺ cells express genes typical of the posterior transcriptional program.

**Figure 4:**
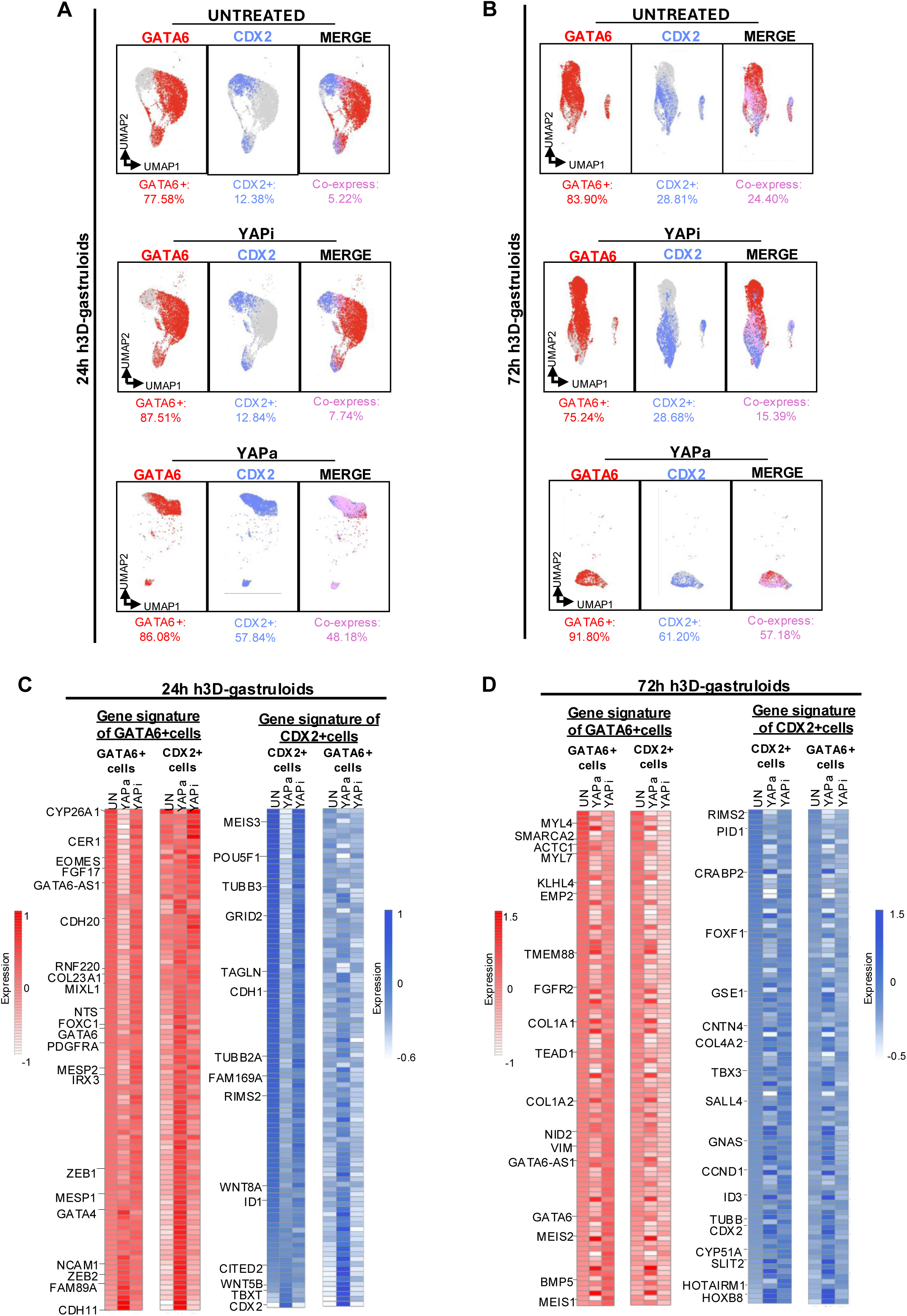
YAP activation compromises the symmetry breaking of 3D gastruloids. (A) UMAP projections of single cell RNA sequencing from 24-hour h3D-gastruloids showing the expression of the lineage markers GATA6 (anterior) and CDX2 (posterior) across untreated, YAPi and YAPa conditions. Quantification of marker expression levels and the percentage of cells co-expressing both markers are shown below each UMAP. (B) Same analysis as in (A) for 72-hour gastruloids. (C) Heatmap depicting expression levels of top 100 expressed genes in GATA6+ cells (“*Gene signature of GATA6+cells”*) of untreated gastruloids, and compared to YAPi and YAPa gastruloids. The expression of these genes in CDX2+ cells, for each condition, is also shown. Heatmaps on the left, show equivalent analysis using the “gene signature of CDX2+cells, at 24 hours. (D) Same as (C) for 72-hour gastruloids.

A reciprocal analysis of the top 100 enriched genes in the GATA6+ cells of the untreated gastruloids (“*GATA6+ cells signature genes*) was performed both at 24 and 72 hours (**Figure 4C-D**). 24-hour YAPa gastruloids revealed broad downregulation of GATA6+ co-expressed genes—along with upregulation of several of these genes within CDX2⁺ cells (e.g., FABP5, LGR5, ZEB2, APLNR, GATA4) (**Figure 4C**). Thus, CDX2⁺ cells in YAPa gastruloids also acquire portions of the anterior gene signature. These findings are fully consistent with the extensive overlap in GATA6 and CDX2 expression observed in **Figure 4A-B**. In contrast, YAPi-treated gastruloids maintained transcriptional profiles closely resembling those of untreated controls, while exhibiting an even sharper and more robust definition of signature gene patterns at 72 hour (**Figure 4C-D**), consistent with an efficient segregation of anterior and posterior domains (**Figure 4B**).

Together, these results demonstrate that forced nuclear YAP activity skews anterior and posterior transcriptional signatures, leading to pervasive co-expression of lineage markers and failure to establish well-defined polarized domains.

## DISCUSSION

Functional studies dissecting the mechanisms that enable hESCs to self-organize into gastruloids remain scarce^8–10,32^. Our findings demonstrate that nuclear retention of the Hippo effector YAP profoundly disrupts gastruloid development by preventing the establishment of a clear anteroposterior axis. In contrast, reducing YAP activity, either through cytoplasmic sequestration or global depletion, promotes robust elongation and the emergence of well-defined anterior and posterior poles. These results reveal YAP levels as a critical regulator of gastruloid development and highlight its role in lineage segregation and axial elongation.

YAP nuclear activity is often associated with proliferative effects, largely through its ability to activate cell cycle regulators^33–39^. However, in our system, YAP activation did not elicit an overt proliferative response, as reflected in the cell-cycle distribution of the scRNA-seq datasets. Instead, YAPa gastruloids exhibited profound disruption of anterior and posterior transcriptional programs, leading to extensive co-expression of lineage markers that are normally restricted to distinct domains. These findings suggest that, in this developmental context, nuclear YAP primarily interferes with lineage specification rather than cell cycle regulation, consistent with our previous work and that of others demonstrating a predominant role for YAP1 in hESC fate decisions^22,25,26,29,40^. Although we did not directly test whether this transcriptional disruption underlies the spherical or ovoid morphology of YAPa gastruloids, impaired lineage segregation is likely a major contributor to their failure to elongate^1,7,41^.

Inhibition of YAP, either chemically or genetically, enhanced gastruloid elongation. Analysis of scRNA-seq datasets suggests that this effect is associated with accelerated differentiation of epiblast-like cells, which were nearly absent in 24-hour YAPi gastruloids. By 72-hour, YAPi gastruloids exhibited an increased proportion of cardiogenic mesoderm and stronger anterior–posterior polarization, as reflected by reduced GATA6/CDX2 co-expression. These findings suggest that limiting YAP activity is necessary for hESCs to undergo timely differentiation into appropriate gastrulation lineages, in agreement with previous observations in other models^22,25,26,29,40^.

Overall, our findings shed light on a novel factor regulating antero-posterior axis elongation in h3D-gastruloids. We and others have previously shown that Wnt, Activin, and BMP4 signaling pathways promote YAP recruitment to the chromatin of developmental genes, including Nodal and Wnt3, in 2D models of differentiation^42^. Thus, it is possible that differential regulation of YAP by critical signaling pathways causes selective YAP binding to target genes across the anteroposterior axis of the gastruloids. We speculate that these selective bindings may be guided by signaling effectors, such as Smads or b-catenin, as shown in other contexts, causing asymmetric regulation of gene expression necessary for the acquisition of distinct lineages. Future investigations will elucidate how developmental signals and YAP cooperate to control the acquisition of lineages along the anteroposterior axis.

### Limitations of the study

Our study combines high-throughput gastruloid generation, single-cell transcriptomics, and complementary chemical and genetic perturbations to uncover the effects of YAP levels on axial elongation in h3D-gastruloids. While these approaches provide strong evidence of the phenotype and differentially expressed genes, we did not distinguish which YAP-regulated genes represent direct versus indirect transcriptional targets, nor did we identify the specific downstream effectors required for elongation. In addition, although the chemical modulators were experimentally validated for their control of YAP activity and closely phenocopy the effects of genetic manipulation of YAP, potential off-target effects cannot be fully excluded.

## METHODS

### Maintenance of hESCs cell culture

WT, YAP KO, doxycycline-inducible YAP (YAP KO:FlagYAP1-PiggyBac) H1 (WAe001-A (RRID:CVCL_9771)) hESCs and YAP KO iPSC cell lines have been described in previous publications from our lab^22,29^. The RUES2 (RUESe002-A (RRID:CVCL_B810)) and RUES2-GLR (RUESe002-A-6) hESC lines were generously provided by Dr. Ali H Brivanlou laboratory and described elsewhere^7,43^. A new YAP KO line was generated in RUES2 hESCs using CRISPR-Cas9 gene editing, following the protocol outlined in^29^. Cell culture methods followed established protocols from our lab^22^. In brief, all cell lines were maintained in incubators set to 37 °C with 5% CO₂. For gastrulation experiments, hESC colonies were maintained in Nutristem hPSC XF medium on Vitronectin-coated plates. Cells were frozen down in a cryopreservation solution containing 10% DMSO and 90% culture media. To minimize spontaneous differentiation, cells were used for no more than five passages before thawing new stocks. For the remaining experiments, a fresh vial was thawed every 20 passages. For ChIP-seq experiments, cells were cultured in mTeSR medium on Matrigel-coated plates. Colonies were passaged at approximately 60–70% confluence using 0.5M PBS/EDTA typically at 1:10 or 1:20 ratio. For single-cell dissociation, Accutase was used to disaggregate the cells, which were then seeded in the presence of ROCK inhibitor for 24 hours to promote survival of single-cells. All cells used in the experiments were tested and confirmed free of mycoplasma and sex specific differences were considered using both male (H1) and female (RUES2) cell lines.

### Generation of human gastruloids using microwell system

The *scale up* protocol for generating human gastruloids was adapted from previously published methods^7^. A microwell system (AggreWell, Stemcell Technologies) was employed following the manufacturer’s guidelines. Briefly, hESC colonies were grown until they reached 40–50% confluency in 6 well-plates and then pre-treated with CHIR99021 at concentrations ranging from 3.25–4 µM (depending on the cell line) in NutriStem medium for 24h. After 24 hours, the cells were dissociated using 0.5 mM EDTA in PBS (-/-) and resuspended in E6 medium containing 1:2000 ROCK inhibitor and CHIR99021 (0.5–1 µM, depending on the cell line). Cells were counted using an automated cell counter. The microwells were prepared by rinsing them with an anti-adherence solution and centrifuging at 2000 × g for 5 minutes at room temperature (RT). The anti-adherence solution was removed, and the cells were seeded at a density of 200–400 cells per microwell in 0.5 mL of E6 medium. Plates were centrifuged at 700 rpm for 2 minutes at RT to facilitate cell aggregation. The medium was replaced daily with fresh E6 for the next two days. Brightfield images were captured using a TIE microscope or EVOS imaging system. To quantify gastruloid elongation, previously described methods were followed^7^. Brightfield images were imported into Fiji software, and the longest axis and midpoint axis of each gastruloid were measured. The elongation ratio was calculated as the ratio of the two measurements and plotted for each condition. To ensure uniformity, the corners of the microwell were avoided, as gastruloids in these areas exhibited more variability (potentially due to inconsistent cell seeding number in the periphery of the well area). To disregard gastruloids in the corners (<10% of total gastruloids in a microwell), we cut the tip of the 200ul pipette, and placed it in the center of the microwell, near to the bottom, while firmly aspirate the gastruloids with accompanied media and transferred them to a new tube for downstream processing. After that, gastruloids in the periphery should remain in the microwell and visible under microscope and can be discarded.

### Whole-mount immunofluorescence of human gastruloids

Staining of the gastruloids was performed and adapted based on previously published literature^44^. h3D-gastruloids were carefully removed from the microwell system using a pipette, transferred through a reversible filter (Stem Cell), and placed in a 6-well dish to eliminate debris. The gastruloids were then fixed in 4% formaldehyde for 25 minutes at room temperature with gentle rocking. Following fixation, the gastruloids were washed three times with PBS. Antigen retrieval was performed with 1% SDS in PBS for 5 minutes and then permeabilization done with 0.1% Triton X-100 in PBS for 30 minutes at RT. After permeabilization, the gastruloids were washed three times with PBS. Primary antibodies (**Table S3**) diluted in PBS with 5% NDS were added, and the gastruloids were incubated overnight at 4°C. The following day, gastruloids were rinsed three times with PBS and then incubated with secondary antibodies diluted in PBS for 1 hour at RT. After secondary incubation, gastruloids were rinsed three times with PBS and then DAPI was added for 10 minutes at RT. Finally, the gastruloids were rinsed three more times with PBS and then mounted onto glass slides, and images were captured using the EVOS M7000 imaging system. Fluorescence intensity quantification was performed using ImageJ.

### Western Blot

Cells were lysed using nuclear lysis buffer (1 M Tris-HCl pH 7, 5 M NaCl, 1 M MgCl2, 10% NP-40, 1 M DTT, and 1x protease inhibitor) on ice for 10 minutes . The lysates were centrifuged at 500 × g for 5 minutes at 4°C, and the supernatant was removed. The pellet was washed with DPBS and protein concentrations were measured using a BCA assay, and each sample was normalized to 25 µg of protein. Proteins were resolved using a pre-cast 4–15% gradient gel, followed by transfer onto a 0.45 µm PVDF membrane using the wet transfer method. The membrane was blocked in 2% BSA in TBST for 45 minutes at room temperature (RT), then incubated overnight at 4°C with the primary antibody. The next day, the membrane was washed three times with TBST for 5 minutes each and incubated with the secondary antibody for 45 minutes at RT. Afterward, the membrane was washed three additional times with TBST (5 minutes per wash). Protein detection was performed using chemiluminescence and imaged on a ChemiDoc system.

### RNA extraction and qRT-PCR

RNA extraction of hESCs was performed using the Quick RNA Extraction Zymo Kit. Between 5 and 10 gastruloids per condition were pooled, washed twice with PBS (-/-), and snap-frozen. Each sample was treated with 700 µL of Trizol, incubated on ice for 2 minutes, followed by the addition of 140 µL of chloroform. The mixture was mixed by hand and incubated at room temperature (RT) for 3 minutes before being centrifuged at 1,000 g for 15 minutes at 4°C. The aqueous phase was carefully transferred to a new tube, and 1/10th of the total volume of 3M sodium acetate was added, along with 1 µL of GlycoBlue. The mixture was gently mixed by hand before adding 350 µL of isopropanol, followed by mixing by hand and incubation at RT for 45 minutes. The samples were then centrifuged at 1,000 g for 45 minutes at 4°C. A visible blue pellet formed at the bottom of the tube, and the supernatant was discarded. The pellet was washed with 700 µL of ethanol, centrifuged at 1,000 g for 10 minutes at 4°C, and the ethanol was removed. Finally, the pellet was resuspended in DNA/RNase-free water, and RNA was quantified using a NanoDrop spectrophotometer. Reverse transcription was performed using 250 ng of RNA with the iScript Reverse Transcription Supermix to generate cDNA. The resulting cDNA was then amplified using the FAST SYBR Green Master Mix on a QuantStudio 3 system. Relative transcript levels were determined using the ΔΔCt method, with RPS23 serving as the normalization control. Primer sequences used for amplification are provided in **Table S4.**

### Sample preparation for single-cell RNAseq analysis of h3D-gastruloids

For single-cell RNA sequencing of h3D-gastruloids, approximately 800-1000 gastruloids from condition were pooled together and filtered through a reversible filter to remove any debris. To ensure uniformity, the corners of the microwell were avoided (as explained above). After, the gastruloids were transferred to a 1.5 mL centrifuge tube and washed with PBS (-/-). Next, the gastruloids were dissociated using TrypLE for 5 minutes at 37°C. After dissociation, the cells were centrifuged, and the pellet was resuspended in media. Cell counting was performed using an automated counter, and cell viability was assessed. Library preparation was carried out using the 10x Chromium Next GEM Single Cell 3’ GEM, Library & Gel Bead Kit v3.1. To assess the quality and concentration of the libraries, Qubit was used for quantification, and Tapestation was employed to check the size distribution. After confirming the library quality, the libraries were pooled together and sequenced on a NextSeq 2000.

### Data analysis of single cell RNAseq

Single-cell sequencing reads were counted individually for each sample using CellRanger (v7.1.0) with default parameters on the hg38 human genome. Doublets and cells contaminated with ambient RNA were assessed using Scrublet and SoupX (v1.6.2) in Python and R, respectively. Low-quality cells, including doublets, cells with high mitochondrial and/or ribosomal content, and those falling in the extreme ends of the RNA feature distribution (very low or very high detected features), were filtered out. Most analyses were performed using the Seurat R package (v4.3.0), including quality control plots generated with the FeatureScatter and VlnPlot functions. Cell cycle phases were assigned to individual cells using the CellCycleScoring function from Seurat, based on expression levels of S and G2/M marker genes. Samples were normalized and variance-stabilized using the SCTransform function in Seurat, regressing out variables such as nCount_RNA, percent.mt, percent.rb, S. Score, and G2M.Score. Principal component analysis (PCA) was run with 30 principal components for all samples. The samples were merged then integrated using Seurat’s SelectIntegrationFeatures, PrepSCTIntegration, FindIntegrationAnchors, and IntegrateData functions with 3000 selected features and the SCTransform normalization method. The RNA assay was used for visualization of gene expression in UMAP and dotplot. Differential gene expression analysis for each cluster between conditions was performed on the RNA assay using the FindMarkers Seurat function after log normalization and scaling. Genes were considered differentially expressed when the adjusted p-value, based on Bonferroni correction, was below 0.05. Cell type count differences between conditions for each cell type were assessed by randomly downsampling the YAPi condition 100 times to match the number of cells in the XMU condition. For each iteration, the number of cells in each cluster was counted. Chi-squared tests with Bonferroni correction were used to assess significant differences in cell counts between genotypes across clusters. Human gastrula scRNAseq data were collected from Tyser et al (2021) imported to Seurat and embedded into a UMAP using the original UMAP coordinates provided by the authors. For the scRNAseq projection, transfer anchors were computed between the human gastrula reference and our gastruloid datasets using PCA-based projection. Our 24h and 72h gastruloids transcriptome were then mapped onto the reference human gastrula UMAP using MapQuery. Prediction scores were extracted from the Seurat transfer anchors and visualized with boxplots to assess the confidence of mapping gastruloid cells onto the human gastrula reference. To assess GATA6/CDX2 co-expression, cells were classified as GATA6+ and/or CDX2+ based on non-zero expression in the Seurat RNA data slot, and single double -positive cells were quantified for each time point. Gene signature of GATA6+ and CDX2+ were identified from the untreated gastruloids condition using Seurat’s FindAllMarkers (min.pct = 0.25, log₂FC > 0.25), and only CDX2 and GATA6 -specific marker genes were kept (ie. genes identified as markers for both CDX2 and GATA6 were removed). For each condition, we then calculated average expression of the top 100 CDX2 and GATA6 -specific signature marker genes within CDX2⁺ or GATA6⁺ subsets and visualized these as heatmaps.

### ChIP-seq and ChIP-qPCR experiments

ChIP was performed as previously described^21^. Briefly, cells were crosslinked with 2 mM DSG for 45 minutes, followed by 1% formaldehyde for 15 minutes. Cells were lysed, sonicated using a Qsonica Q700, and non-sonicated material was removed by centrifugation. For immunoprecipitation, 300 µL of chromatin extract was incubated with 1–5 µg of primary antibody overnight at 4°C. Protein A/G magnetic beads (10 µL) were added, and incubation continued for 4 hours at 4°C. The complexes were washed four times with buffers listed in **Table S4**. Crosslinking was reversed by incubating overnight at 65°C in elution buffer (1% SDS, 0.1 M NaHCO3). DNA was purified using the QIAquick PCR purification kit. For ChIP-qPCR, DNA was amplified with FAST SYBR Green Master Mix on a Quant Studio 3, using 2 µL of eluted DNA per PCR reaction. DNA concentration was measured with a Qubit Flex, and libraries were prepared using the NEBNext DNA Library Kit. Samples were sequenced on a NextSeq 1000. Primers and antibodies are listed in **Table S4**.

### Statistical Methods

Statistical analysis was conducted to characterize gastruloids and evaluate differences between experimental groups. The statistical test performed depended on the comparison and the number of groups analyzed. For comparisons between two groups, a two-tailed Student’s t-test was used and for comparisons involving more than two groups, a one-way ANOVA was used. For scRNAseq experiments, chi-squared tests with Bonferroni correction was used to analyze cell type distribution and proportions. Each experiment was conducted with three biological replicates unless otherwise specified. For gastruloid experiments, approximately 10-50 gastruloids were generated per experimental condition. All statistical analyses were performed using GraphPad Prism and R.

## RESOURCE AVAILABILITY

### LEAD CONTACT

Requests for further information and resources should be directed to the led contact, Conchi Estaras.

## MATERIAL AVAILABILITY

The cell lines used in this study are available from public repositories.

## ACKNOWLEDGMENTS

We acknowledge all current and previous members of the Estaras lab for discussions, feedback, and assistance in experiments. Also, thanks to Dr. Elrod, Dr. Simon, Dr. Kishore and Dr. Garikipati’s laboratories at Aging+Cardiovascular Discovery Center for discussions and feedback. Thanks to Dr. Kishore’s lab for their generosity sharing reagents. We thank Dr. Johnathan Whetstine, their laboratory members Madison Honer, Dr. Zach Gray, and Ben Ferman, and the Genomics Facility at the FCCC, for sharing reagents and for the single-cell training. This work was funded by the NIH/NICHD R01 HD106969 (to C.E), NIH/NHLBI 5T32HL091804 (to E.A), NIH/NICHD F31HD113419 (to E.A).

## AUTHORS CONTRIBUTIONS

Conceptualization, C.E., E.A.; Data curation, C.E., E.A., T.R., and N.A.; Formal analysis, E.A., T.R., and N.A.; Funding acquisition, C.E. and E.A.; Investigation, C.E., E.A., T.R., O.M.P, and M.Z.; Methodology, E.A.; Supervision, C.E.; Visualization, E.A., T.R., and N.A.; Writing – original draft, C.E. and E.A.; Writing- reviewing and editing, C.E. and E.A

## REFERENCES

1 Ghimire, S., Mantziou, V., Moris, N. & Arias, A. M. Human gastrulation: The embryo and its models. Dev Biol (2021). 10.1016/j.ydbio.2021.01.006

2 Wolpert, L. Gastrulation and the evolution of development. Dev Suppl, 7–13 (1992).

3 Zhai, J., Xiao, Z., Wang, Y. & Wang, H. Human embryonic development: from peri-implantation to gastrulation. Trends Cell Biol (2021). 10.1016/j.tcb.2021.07.008

4 Ferrer-Vaquer, A. & Hadjantonakis, A. K. Birth defects associated with perturbations in preimplantation, gastrulation, and axis extension: from conjoined twinning to caudal dysgenesis. Wiley Interdiscip Rev Dev Biol 2, 427–442 (2013). 10.1002/wdev.97

5 Stutt, N., Song, M., Wilson, M. D. & Scott, I. C. Cardiac specification during gastrulation - The Yellow Brick Road leading to Tinman. Semin Cell Dev Biol 127, 46–58 (2022). 10.1016/j.semcdb.2021.11.011

6 Herion, N. J., Salbaum, J. M. & Kappen, C. Traffic jam in the primitive streak: the role of defective mesoderm migration in birth defects. Birth Defects Res A Clin Mol Teratol 100, 608–622 (2014). 10.1002/bdra.23283

7 Moris, N. et al. An in vitro model of early anteroposterior organization during human development. Nature 582, 410–415 (2020). 10.1038/s41586-020-2383-9

8 Dingare, C., Cao, D., Yang, J. J., Sozen, B. & Steventon, B. Mannose controls mesoderm specification and symmetry breaking in mouse gastruloids. Dev Cell 59, 1523–1537.e1526 (2024). 10.1016/j.devcel.2024.03.031

9 Underhill, E. J. & Toettcher, J. E. Control of gastruloid patterning and morphogenesis by the Erk and Akt signaling pathways. Development 150 (2023). 10.1242/dev.201663

10 McNamara, H. M., Solley, S. C., Adamson, B., Chan, M. M. & Toettcher, J. E. Recording morphogen signals reveals mechanisms underlying gastruloid symmetry breaking. Nat Cell Biol 26, 1832–1844 (2024). 10.1038/s41556-024-01521-9

11 Rossi, G., Giger, S., Hübscher, T. & Lutolf, M. P. Gastruloids as in vitro models of embryonic blood development with spatial and temporal resolution. Sci Rep 12, 13380 (2022). 10.1038/s41598-022-17265-1

12 Beccari, L. et al. Multi-axial self-organization properties of mouse embryonic stem cells into gastruloids. Nature 562, 272–276 (2018). 10.1038/s41586-018-0578-0

13 Girgin, M. U., Broguiere, N., Mattolini, L. & Lutolf, M. P. Gastruloids generated without exogenous Wnt activation develop anterior neural tissues. Stem Cell Reports 16, 1143–1155 (2021). 10.1016/j.stemcr.2021.03.017

14 Shahbazi, M. N., Siggia, E. D. & Zernicka-Goetz, M. Self-organization of stem cells into embryos: A window on early mammalian development. Science 364, 948–951 (2019). 10.1126/science.aax0164

15 Vincent, S. D., Dunn, N. R., Hayashi, S., Norris, D. P. & Robertson, E. J. Cell fate decisions within the mouse organizer are governed by graded Nodal signals. Genes Dev 17, 1646–1662 (2003). 10.1101/gad.1100503

16 Camus, A., Perea-Gomez, A., Moreau, A. & Collignon, J. Absence of Nodal signaling promotes precocious neural differentiation in the mouse embryo. Dev Biol 295, 743–755 (2006). 10.1016/j.ydbio.2006.03.047

17 Robertson, E. J. Dose-dependent Nodal/Smad signals pattern the early mouse embryo. Semin Cell Dev Biol 32, 73–79 (2014). 10.1016/j.semcdb.2014.03.028

18 Kowalczyk, W. et al. Hippo signaling instructs ectopic but not normal organ growth. Science 378, eabg3679 (2022). 10.1126/science.abg3679

19 Rito, T. et al. Timely TGFβ signalling inhibition induces notochord. Nature (2024). 10.1038/s41586-024-08332-w

20 Morin-Kensicki, E. M. et al. Defects in yolk sac vasculogenesis, chorioallantoic fusion, and embryonic axis elongation in mice with targeted disruption of Yap65. Mol Cell Biol 26, 77–87 (2006). 10.1128/MCB.26.1.77-87.2006

21 Wu, Z. & Guan, K. L. Hippo Signaling in Embryogenesis and Development. Trends Biochem Sci 46, 51–63 (2021). 10.1016/j.tibs.2020.08.008

22 Stronati, E., et al. YAP1 regulates the self-organized fate patterning of hESC-derived gastruloids. Stem Cell Reports (2022). 10.1016/j.stemcr.2021.12.012

23 Beyer, T. A. et al. Switch enhancers interpret TGF-β and Hippo signaling to control cell fate in human embryonic stem cells. Cell Rep 5, 1611–1624 (2013). 10.1016/j.celrep.2013.11.021

24 Tyser, R. C. V. et al. Single-cell transcriptomic characterization of a gastrulating human embryo. Nature 600, 285–289 (2021). 10.1038/s41586-021-04158-y

25 Abraham, E. et al. A retinoic acid:YAP1 signaling axis controls atrial lineage commitment. Cell Rep 44, 115687 (2025). 10.1016/j.celrep.2025.115687

26 Abraham, E. et al. YAP1 and QSER1 are Key Modulators of Embryonic Signaling Pathways in the Mammalian Epiblast. bioRxiv (2025). 10.1101/2025.06.16.659935

27 Fan, F. et al. Pharmacological targeting of kinases MST1 and MST2 augments tissue repair and regeneration. Sci Transl Med 8, 352ra108 (2016). 10.1126/scitranslmed.aaf2304

28 Tang, Z. et al. A brief review: some compounds targeting YAP against malignancies. Future Oncol 15, 1535–1543 (2019). 10.2217/fon-2019-0035

29 Estarás, C., Hsu, H. T., Huang, L. & Jones, K. A. YAP repression of the WNT3 gene controls hESC differentiation along the cardiac mesoderm lineage. Genes Dev 31, 2250–2263 (2017). 10.1101/gad.307512.117

30 Xin, M. et al. Regulation of insulin-like growth factor signaling by Yap governs cardiomyocyte proliferation and embryonic heart size. Sci Signal 4, ra70 (2011). 10.1126/scisignal.2002278

31 De Santis, R. et al. Crosstalk between tissue mechanics and BMP4 signaling regulates symmetry breaking in human gastrula models. Cell Stem Cell 32, 1691–1704.e1696 (2025). 10.1016/j.stem.2025.09.006

32 Mantziou, V. et al. In vitro teratogenicity testing using a 3D, embryo-like gastruloid system. Reprod Toxicol 105, 72–90 (2021). 10.1016/j.reprotox.2021.08.003

33 Lee, H. C. et al. YAP1 overexpression contributes to the development of enzalutamide resistance by induction of cancer stemness and lipid metabolism in prostate cancer. Oncogene 40, 2407–2421 (2021). 10.1038/s41388-021-01718-4

34 LeBlanc, L., Ramirez, N. & Kim, J. Context-dependent roles of YAP/TAZ in stem cell fates and cancer. Cell Mol Life Sci 78, 4201–4219 (2021). 10.1007/s00018-021-03781-2

35 Kim, H. R., Seo, C. W., Yoo, K., Han, S. J. & Kim, J. Yes-associated protein 1 as a prognostic biomarker and its correlation with telomerase in various cancers. Osong Public Health Res Perspect 12, 324–332 (2021). 10.24171/j.phrp.2021.0207

36 Barry, E. R. et al. Restriction of intestinal stem cell expansion and the regenerative response by YAP. Nature 493, 106–110 (2013). 10.1038/nature11693

37 von Gise, A. et al. YAP1, the nuclear target of Hippo signaling, stimulates heart growth through cardiomyocyte proliferation but not hypertrophy. Proc Natl Acad Sci U S A 109, 2394–2399 (2012). 10.1073/pnas.1116136109

38 Rosenbluh, J. et al. β-Catenin-driven cancers require a YAP1 transcriptional complex for survival and tumorigenesis. Cell 151, 1457–1473 (2012). 10.1016/j.cell.2012.11.026

39 Schlegelmilch, K. et al. Yap1 acts downstream of α-catenin to control epidermal proliferation. Cell 144, 782–795 (2011). 10.1016/j.cell.2011.02.031

40 Hsu, H. T., Estarás, C., Huang, L. & Jones, K. A. Specifying the Anterior Primitive Streak by Modulating YAP1 Levels in Human Pluripotent Stem Cells. Stem Cell Reports 11, 1357–1364 (2018). 10.1016/j.stemcr.2018.10.013

41 van den Brink, S. C. & van Oudenaarden, A. 3D gastruloids: a novel frontier in stem cell-based in vitro modeling of mammalian gastrulation. Trends Cell Biol 31, 747–759 (2021). 10.1016/j.tcb.2021.06.007

42 Stronati, E. et al. YAP1 regulates the self-organized fate patterning of hESC-derived gastruloids. Stem Cell Reports 17, 211–220 (2022). 10.1016/j.stemcr.2021.12.012

43 Martyn, I., Kanno, T. Y., Ruzo, A., Siggia, E. D. & Brivanlou, A. H. Self-organization of a human organizer by combined Wnt and Nodal signalling. Nature 558, 132–135 (2018). 10.1038/s41586-018-0150-y

44 Baillie-Johnson, P., van den Brink, S. C., Balayo, T., Turner, D. A. & Martinez Arias, A. Generation of Aggregates of Mouse Embryonic Stem Cells that Show Symmetry Breaking, Polarization and Emergent Collective Behaviour In Vitro. J Vis Exp (2015). 10.3791/53252

45 Allhoff, M., Seré, K., F Pires, J., Zenke, M. & G Costa, I. Differential peak calling of ChIP-seq signals with replicates with THOR. Nucleic Acids Res 44, e153 (2016). 10.1093/nar/gkw680

