## Supplemental Figures for "YAP levels regulate anteroposterior elongation of hESC-derived gastruloids"

**Supplementary Figure 1. Supports Figure 1.**

(A) Representative whole-mount immunostaining of 72-hour RUES2-derived h3D-gastruloids showing polarized expression of CDX2 (posterior marker) and GATA6 (anterior marker). Scale bar, 200 $\mu m$. The experiment was performed twice with similar results.

(B-C) UMAP projection of single-cell RNA seqeucning  from 24 (B) and 72-hour h3D-gastruloids (C) showing expression of key developmental lineage markers used to define major populations: EPI, epiblast; PS, primitive streak; ENDO, endoderm; HEMTAO: hematopoietic

(D and E) Prediction scores mapping 24 (D) and 72-hour (E) 3D-gastruloid cell clusters onto the CS7 human gastrula reference atlas from Tyser *et al*., 2021, demonstrating similarity between in vitro–derived populations and in vivo human gastrulation states.

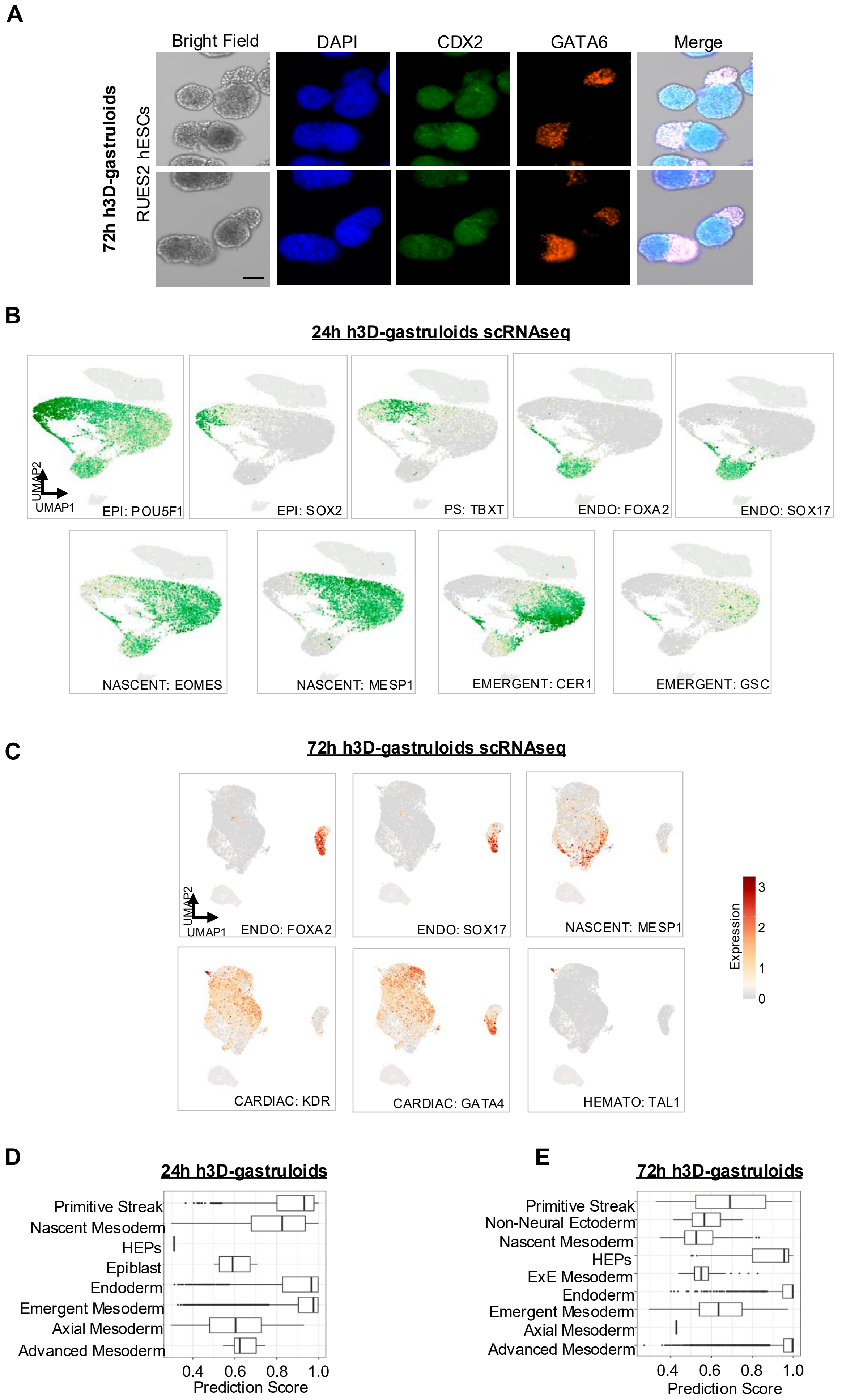
A
Bright Field
DAPI
CDX2
GATA6
Merge
72h h3D-gastruloids
RUES2 hESCs
B
24h h3D-gastruloids scRNAseq
UMAP2
UMAP1
EPI: POU5F1
EPI: SOX2
PS: TBXT
ENDO: FOXA2
ENDO: SOX17
NASCENT: EOMES
NASCENT: MESP1
EMERGENT: CER1
EMERGENT: GSC
Expression
0
1
2
3
C
72h h3D-gastruloids scRNAseq
UMAP2
UMAP1
ENDO: FOXA2
ENDO: SOX17
NASCENT: MESP1
CARDIAC: KDR
CARDIAC: GATA4
HEMATO: TAL1
Expression
3
2
1
0
D
24h h3D-gastruloids
Primitive Streak
Nascent Mesoderm
HEPs
Epiblast
Endoderm
Emergent Mesoderm
Axial Mesoderm
Advanced Mesoderm
0.4
0.6
0.8
1.0
Prediction Score
E
72h h3D-gastruloids
Primitive Streak
Non-Neural Ectoderm
Nascent Mesoderm
HEPs
ExE Mesoderm
Endoderm
Emergent Mesoderm
Axial Mesoderm
Advanced Mesoderm
0.4
0.6
0.8
1.0
Prediction Score

**Supplementary Figure 2. Supports Figure 2.**

(A) qPCR analysis of canonical Hippo/YAP targert genes CTGF and CYR61 in WT hESCs treated with the indicated conditions for 48 hours. Data are shown as mean ± SEM. Experiment was performed once with three technical replicates. Statistical analysis: one-way ANOVA (Kruskal–Wallis test). $p < 0.05$ (*), $p < 0.01$ (**).

(B) Full Western blot image of YAPa- and YAPi- treated nuclear WT H1 hESCs. Red arrows indicate regions used for the cropped blots shown in the main figure.

(C) Representative bright-field images of 48-hour RUES2-derived h3D-gastruloids untreated or treated with YAP inhibitor or YAP activator. Scale bar, 125um. Quantifications corresponding to these images are presented in the main figure.

(D) Western blot analysis of YAP protein levels in WT and YAP KO RUES2 and H1 hESC lines generated using CRISPR/Cas9. GAPDH was used as a loading control.

(E) Immunostaining of NANOG and SOX2 in WT and YAP KO RUES2 hESCs. Scale bar, 50um.

(F) Fluorescence intensities of SOX2 and NANOG were quantified and normalized to DAPI. The experiment was performed once with three technical replicates. A total of 30 cells were analyzed (10 cells per image across three images). Statistical analysis: Student's *t*-test.

(G) Representative images of 48 and 72-hour WT and YAP KO h3D-gastruloids derived from multiple genetic backgrounds (RUES2, H1, and iPSCs). Scale bar, 200 µm. Quantifications are shown in the main figure.

#### Supplementary Figure 2

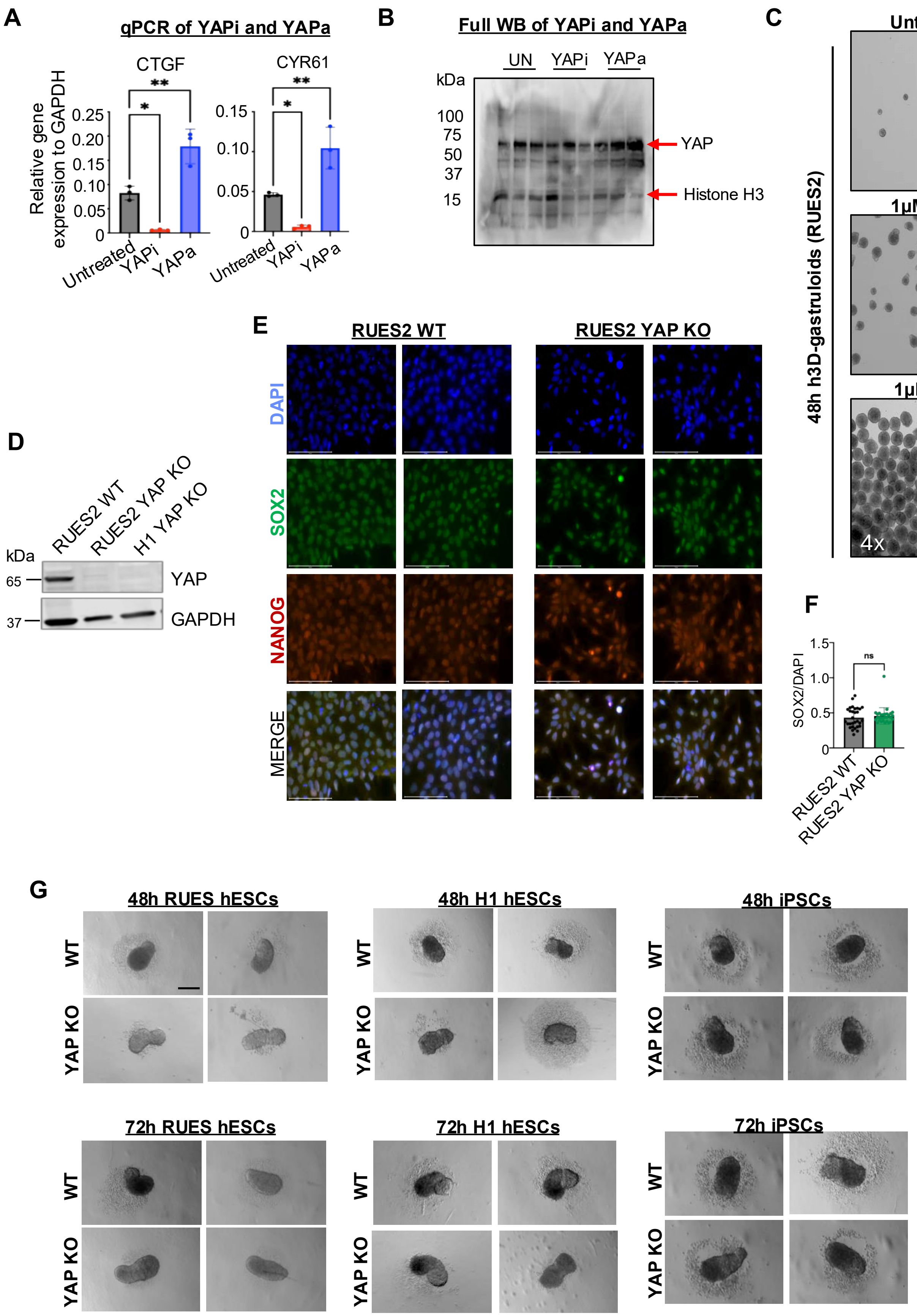

**Supplementary Figure 3. Supports Figure 3.**

(A) UMAP projections showing the expression of defined YAP-target genes for 24 and 72-hour h3D-gastruloids, demonstrating the effectiveness of YAP inhibitor (YAPi) and YAP activator (YAPa) treatments

(B) Cell cycle scoring (G1, S, and G2/M phases) derived from scRNAseq of 24 and 72-hour gastruloids treated with YAPa or YAPi.

(C) UMAP [projections showing the expression of key lineage-defining markers in 24-hour h3D-gastruloids.

### Supplementary Figure 3

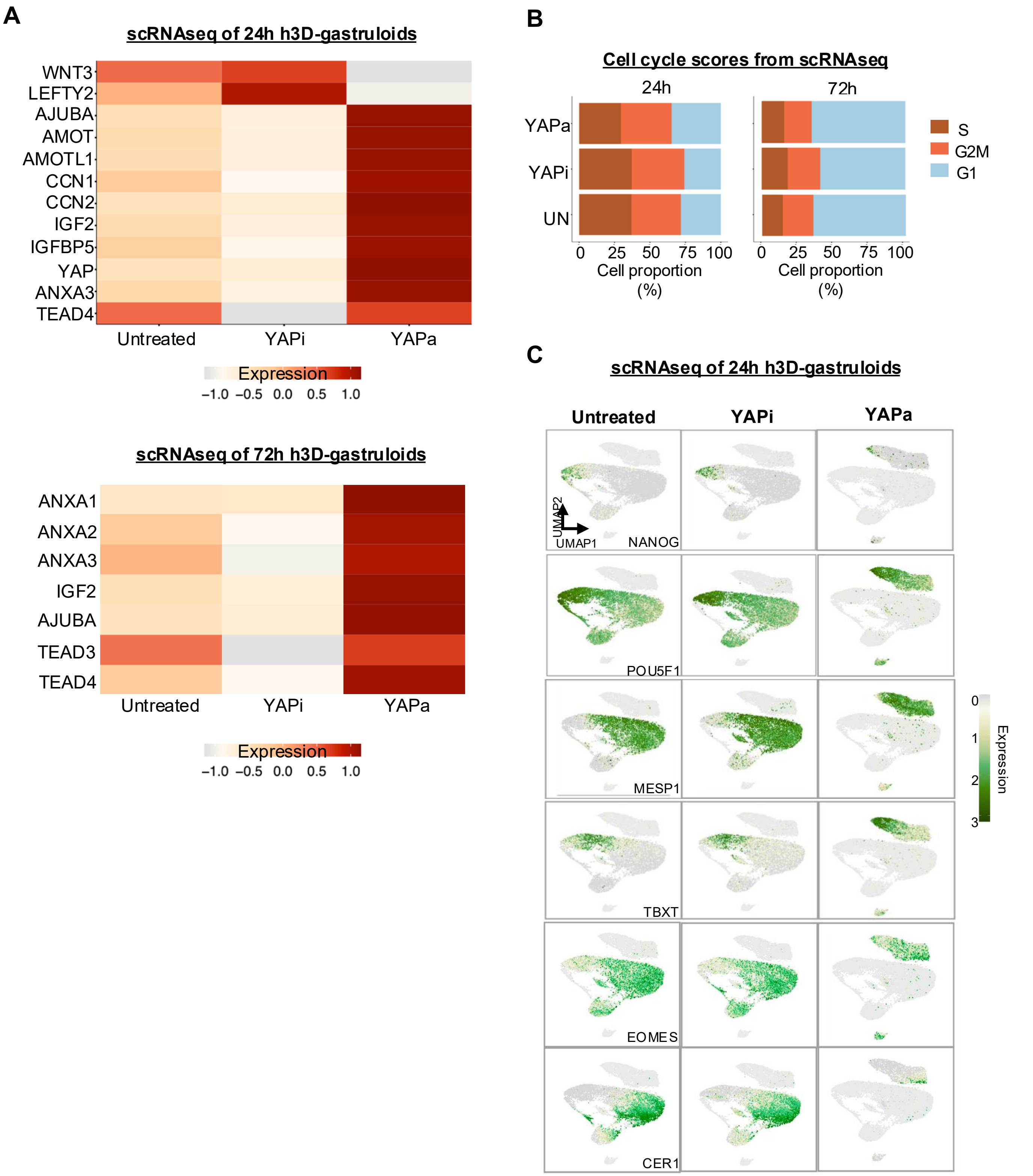

**Supplementary Figure 4. Supports Figure 4.**

(A) Whole-mount immunostaining of CDX2 (posterior marker) and GATA6 (anterior marker) in untreated72-hour gastruloids. Scale bar, *50 µm.*

(B) Untreated and YAP activator (YAPa)-treated 72-hour h3D-gastruloids were dissociated and immunostained for YAP, CDX2, and GATA6. Fluorescence intensity of CDX2 and GATA6 was normalized to DAPI. Experiments were performed once with 6–10 gastruloids per condition, and 30 cells were quantified per condition. Scale bar, 100 µm. Statistical analysis: Student's t-test ****p < 0.0001.

### Supplementary Figure 4

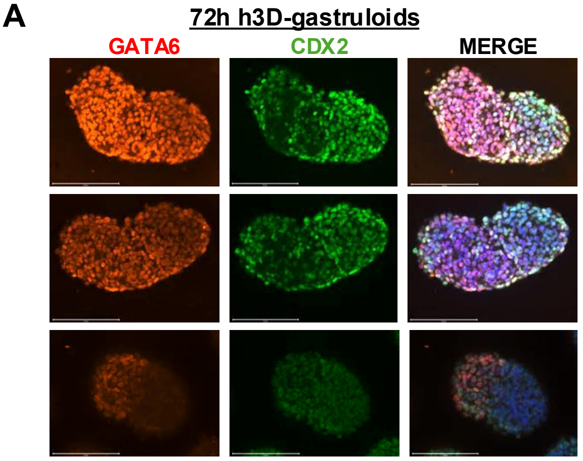


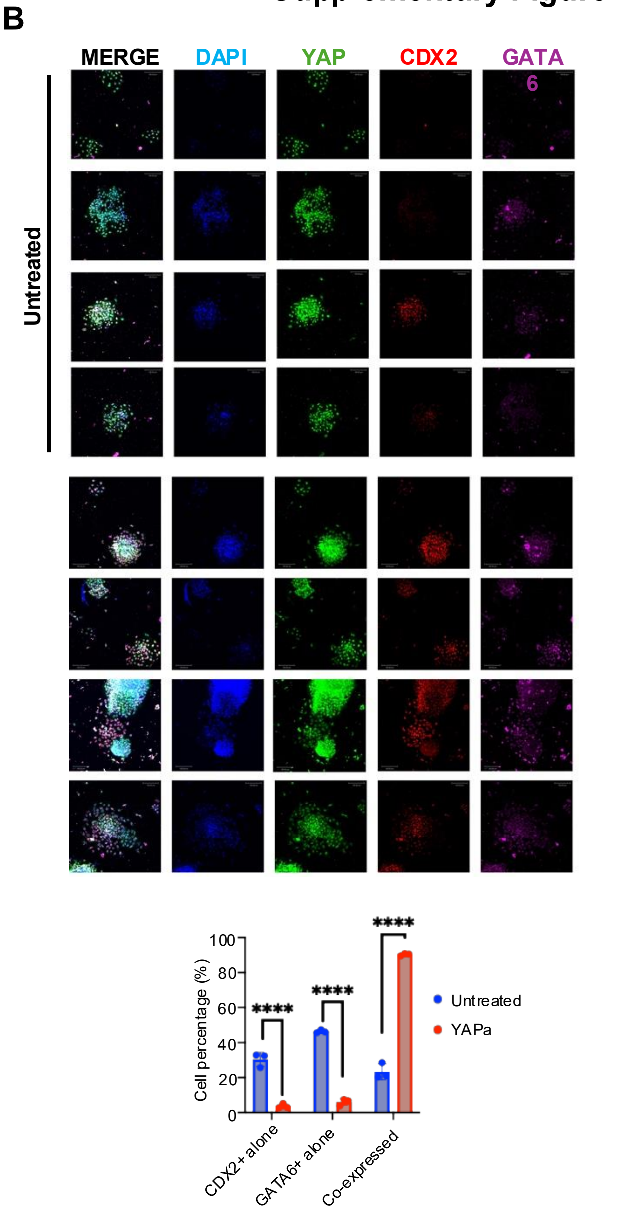
